# A microbial mutualist within host individuals increases parasite transmission between host individuals: Evidence from a field mesocosm experiment

**DOI:** 10.1101/2021.11.28.470289

**Authors:** Kayleigh R. O’Keeffe, Brandon T. Wheeler, Charles E. Mitchell

**Affiliations:** Department of Biology, University of North Carolina at Chapel Hill, Chapel Hill, NC, USA; Department of Biology, University of Pennsylvania, Philadelphia, PA, USA; Department of Biology, Western Carolina University, Cullowhee, NC, USA; Environment, Ecology and Energy Program, University of North Carolina, Chapel Hill, NC, USA

**Author notes:** Correspondence Kayleigh O’Keeffe.

**Keywords:** within-host microbial interactions, species interactions, co-infection, mesocosm experiment, transmission, disease ecology, plant pathogens

## Abstract

The interactions among host-associated microbes and parasites can have clear consequences for disease susceptibility and progression within host individuals. Yet, empirical evidence for how these interactions impact parasite transmission between host individuals remains scarce. We address this scarcity by using a field mesocosm experiment to investigate the interaction between a systemic fungal endophyte, *Epichloë coenophiala*, and a fungal parasite, *Rhizoctonia solani*, in leaves of a grass host, tall fescue. Specifically, we investigated how this interaction impacted parasite transmission under field conditions in replicated experimental host populations. *Epichloë*-inoculated populations tended to have greater disease prevalence over time, though this difference had weak statistical support. More clearly, *Epichloë*-inoculated populations experienced higher peak parasite prevalences than *Epichloë*-free populations. *Epichloë* conferred a benefit in growth; *Epichloë*-inoculated populations had greater aboveground biomass than *Epichloë*-free populations. Using biomass as a proxy, host density was correlated with peak parasite prevalence, but *Epichloë* still increased peak parasite prevalence after controlling for the effect of biomass. Together, these results suggest that within-host microbial interactions can impact disease at the population level. Further, while *Epichloë* is clearly a mutualist of tall fescue, it may not be a defensive mutualist in relation to *R. solani*.

## Introduction

Parasites commonly interact with other parasites, as well as commensals and mutualists, within a shared host individual (Borer, Kinkel, May, & Seabloom, 2013; Tollenaere, Susi, & Laine, 2016). Host organisms are commonly infected by defensive symbionts, which can interact with parasites to protect their hosts (reviewed in K. Clay, 2014; Hopkins, Wojdak, & Belden, 2017). These defensive symbionts can have impacts on disease at the individual and population-levels (Hopkins et al., 2017; O’Keeffe, Carbone, Jones, & Mitchell, 2017). As the ubiquity of diverse within-host microbial communities have come to be realized, a challenge has been to link within- and between-host levels of disease dynamics (P. A. Clay, Duffy, & Rudolf, 2020). Here, we address this challenge by investigating the within- and between-host impacts of a defensive symbiont of a grass host on the severity and spread of a fungal parasite under field conditions.

Defensive symbionts can dramatically impact the survivorship, growth, and reproduction of parasites infecting the same host individual (Arnold et al., 2003; Costello, Stagaman, Dethlefsen, Bohannan, & Relman, 2012; Santhanam et al., 2015). Within a host, defensive symbionts may prime a host immune response to parasites (Selosse, Bessis, & Pozo, 2014) or directly interfere with an invading parasite (Gerardo & Parker, 2014). For example, systemic fungal endophytes of grasses can produce antimicrobial compounds that may reduce severity of disease caused by a parasite on a plant individual (K. Clay, Cheplick, & Marks, 1989). Through such mechanisms, defensive symbionts can affect host susceptibility, parasite growth and replication, and subsequent parasite disease severity (Arnold et al., 2003; Oliver, Moran, & Hunter, 2005; Hussain et al., 2013).

Interactions among coinfecting symbionts within host individuals can have implications that scale up to populations (Cattadori, Boag, & Hudson, 2008; Telfer et al., 2010). Defensive symbionts can impact the growth and reproduction of parasites within a host and within-host accumulation is often directly or indirectly linked to between-host transmission (Wintermantel, Cortez, Anchieta, Gulati-Sakhuja, & Hladky, 2008; Tollenaere et al., 2016). Defensive symbionts may therefore be an important driver of epidemiological dynamics, which can have impacts on ecosystem function (Paseka et al., 2020). Yet, how within-host dynamics of defensive symbionts and parasites scale to impacts across host individuals remains an important frontier in disease ecology (Viney & Graham, 2013; Ezenwa & Jolles, 2015; Johnson, de Roode, & Fenton, 2015).

To investigate the impacts of a defensive symbiont on a parasite across levels of ecological organization, we conducted a field mesocosm experiment on a vertically transmitted fungal endophyte, *Epichloë coenophiala*, the facultative fungal parasite *Rhizoctonia solani*, and a host grass, tall fescue (*Lolium arundinaceum*). We established experimental populations of *Epichloë*-inoculated and *Epichloë*-free plants in field mesocosms, inoculated them with the parasite, then performed repeated surveys of parasite damage on leaves. We provide evidence that while this endophyte clearly has a mutualistic relationship with its host at the host-individual level, it can have a contrasting impact on parasite transmission at the host-population level.

## Methods

### Study System

Within a grass host, tall fescue (*Lolium arundinaceum*), we investigated the interaction between two fungal symbionts: the parasite *Rhizoctonia solani* and the vertically-transmitted systemic endophyte *Epichloë coenophiala. R. solani* is a facultative parasite, as it can persist in the soil as a saprobe. As a necrotrophic parasite, it extracts resources by killing host cells. In contrast, *E. coenophiala* is considered a mutualist under most ecological conditions (Kari Saikkonen, Young, Helander, & Schardl, 2016). While *E. coenophiala* consistently acts as a defensive mutualist with regards to herbivory, *E. coenophiala* can vary in its impact on parasites (Potter, 1980, 1982; Liu, Kang, & Buchenauer, 2006; Kari Saikkonen, Gundel, & Helander, 2013). Empirical evidence for the direction of the interaction between *Epichloë* endophytes and *Rhizoctonia* parasites varies (Pańka, West, Guerber, & Richardson, 2013; Halliday, Umbanhowar, & Mitchell, 2017; O’Keeffe, Simha, & Mitchell, 2021)

### Experimental Design and Setup

We investigated how within-host microbial interactions impact parasite transmission by conducting two field mesocosm experiments. This experiment was conducted at Widener Farm, an old field of the Duke Forest Teaching and Research Laboratory (Orange County, NC, USA), during the summer of 2018. This old field produced row crops until 1996 and has since been mowed to produce hay. During the 2013-2017 growing seasons, surveys of the tall fescue population at this site showed that symptoms from parasite, *Rhizoctonia solani*, began appearing on leaves in June or July, peaked in prevalence in August and September, and decreased in prevalence over the fall months (Halliday, Umbanhowar, & Mitchell, 2017). Because the peak of this parasite epidemic at this site occurred in August in previous years (Halliday et al., 2017), we conducted each experiment during that time period in subsequent years, 2017 and 2018. While the overall design of each experiment was similar in 2017 and 2018, there were a few key differences and notably, parasite transmission was more successful in the 2018 experiment (Figure S1). Owing to the relative lack of transmission in 2017, we report the 2017 transmission methods and results in the supplement (supplementary methods and Table S1), and here in the main text, we report the 2018 methods and results.

To test how the endophyte affects parasite spread across a host population, we manipulated presence of *E. coenophiala* at the level of the host population. We planted a total of 26 populations, and each population was contained within a 45-inch (1.14 meter) diameter plastic wading pool (Summer Escapes) to limit *R. solani* inoculum coming from the environment. Each population was randomly assigned one of two treatments: *E. coenophiala*-inoculated or *E. coenophiala*-free (herein referred to has *Epichloë*-inoculated and *Epichloë*-free). In total, there were 15 *Epichloë*-inoculated and 11 *Epichloë*-free populations. Two randomly selected populations in each *Epichloë* treatment (4 total) were not inoculated with the parasite and served as experimental controls.

Each population consisted of 13 plants: 1 plant in the center of the population (which would ultimately be inoculated with the parasite), and 12 plants surrounding the central plant at distances of 12cm, 24cm, and 36cm away (4 plants at each distance, Figure S2). The 338 plants within the experiment were propagated from *Epichloë*-free or *Epichloë*-inoculated seed produced by Tim Phillips at the University of Kentucky and the Noble Research Institute in Ardmore Oklahoma, respectively. All seed was from variety KY-31. Seed was germinated on June 25, 2018 and grown in a greenhouse for 6 weeks. The greenhouse temperature was kept between 19.7-22.2°C and light was supplemented between 9am and 7pm if natural light fell below 350 W/m^2^. All plants except for the central plants were transplanted into the contained field mesocosm experiment on Monday, August 6, 2018, 6 weeks after germination. Each population consisted of plants belonging to the same endophyte category (all *Epichloë*-inoculated or *Epichloë*-free) except all central plants were *Epichloë*-free. Plants were randomly assigned to one of the populations, and locations of the plants within the populations were also randomized. The populations were fully randomized in a 2 × 13 layout, with narrow paths separating populations (Figure S2). The plants were given four days to acclimate to the field prior to the introduction of the parasite.

The plants that would ultimately be planted in the central position of the populations were transferred to growth chambers on August 8, 2018 and inoculated with an isolate of the parasite that was cultured in 2015 from a leaf lesion on a tall fescue plant in the same field as this experiment. Once in axenic culture, plugs of the leading edge of the culture were stored in mineral oil and potato dextrose broth in a -80°C freezer. We plated these plugs on potato dextrose agar and the resulting growth served as the source of inoculum for this experiment. Inoculum consisted of a 6mm diameter plug of potato dextrose agar with the leading edge of the parasite culture placed directly at the base of a leaf. Parasite infection success depends on a humid environment. To maintain moisture at the site of inoculation, a cotton ball wet with sterile water was placed over the inoculum, secured with tin foil and parafilm. The inoculated plants were placed in a growth chamber (Percival PGC-6L (Perry, Iowa)) for two days with a 12 hour light/12 hour dark cycle set at 28°C and humidity was maintained at approximately 95% with humidifiers (Vicks V5100-N Ultrasonic Humidifier) on each shelf of the growth chambers. In addition to parasite-inoculated plants, four plants were mock-inoculated with plugs of potato dextrose agar without *R. solani* mycelia. After two days, all plants inoculated with *R. solani* exhibited parasite symptoms and were transplanted into the field mesocosm experiment on August 10, 2018. One mock-inoculated plant was transplanted into each of the four experimental control populations.

### Data Collection

Following the placement of the central inoculated plant, twice a week, for four weeks, 7 random leaves per plant were non-destructively selected and observed for the presence or absence of damage caused by the parasite, as well as any other parasite damage. Specifically, leaves were surveyed 4, 8, 11, 14, 18, 22, 25, and 28 days after parasite inoculation (8 surveys total). Each leaf was surveyed for the presence of any damage caused by parasites, herbivores, or abiotic sources.

To measure disease severity over time, percent leaf area damaged by the parasite was quantified on individual leaves on one tagged tiller per plant once a week. On each leaf, the initial date of symptomatic infection by the parasite was recorded, and severity was estimated by visually comparing leaves to reference images of leaves of known infection severity (Mitchell, Tilman, & Groth, 2002, 2003; Halliday et al., 2017). Over the course of the experiment, three severity surveys were conducted (8, 14, and 22 days after parasite inoculation).

At the conclusion of the experiment, we collected and froze (−20°C freezer) inch-long cross-sections of two tillers per plant to confirm endophyte presence. Endophyte infection was tested via immunoblot (Agrinostics Ltd. Co, Watkinsville, GA). Additionally, aboveground biomass was harvested, dried and weighed.

### Data Analysis

*Epichloë*-inoculated seed did not always result in endophyte detection in aboveground tissue. Overall, we detected the endophyte in aboveground tissue in 42% of *Epichloë*-inoculated plants. This resulted in variation in endophyte prevalence among the *Epichloë*-inoculated populations (minimum: 15.4%; maximum: 69.2%). We therefore analyzed our data in two separate ways: with endophyte treatment (2 levels: *Epichloë*-free or *Epichloë*-inoculated) as a predictor, or with endophyte prevalence (continuous variable from 0-1) as a predictor.

Control populations (in which the central plant was mock-inoculated) did not exhibit symptoms of the parasite, confirming that mesocosm populations were not infected from environmental sources of inoculum. These control populations were therefore excluded from analyses.

The models based on parasite prevalence reported in the main text only include surveys until each population’s peak parasite prevalence because we were interested in how endophyte presence impacts epidemic spread, which is no longer happening after the peak. We report the results of models based on parasite prevalence with all surveys in the supplement (Table S2, Figure S3). All analyses were performed in R, version 3.6.1.

Leaves were analyzed as host individuals because each parasite infection is restricted to a single leaf (as done by Halliday et al., 2017; O’Keeffe et al., 2021). To summarize disease progression over time, area under the disease progress stairs (AUDPS) was calculated for each population using the audps function within the agricolae package (version 1.3, De Mendiburu 2016). AUDPS estimated the integration of the development of disease progress experienced by each population by adding together polygon steps between each time point (Simko & Piepho, 2011). Disease intensity at each time point was calculated separately with two types of data: data from the weekly severity surveys, represented by the average leaf area damaged across all leaves surveyed within a population and data from the twice-weekly prevalence surveys, represented by the parasite prevalence. When calculated with prevalence data, AUDPS was log transformed to achieve homoscedasticity and normality of residuals. We investigated if and how endophyte treatment affected these estimates of disease progression over time with a linear model.

To further evaluate the magnitude of epidemics, we investigated measurements of parasite prevalence repeated over time using linear mixed effects models. Data on proportion of leaves infected with the parasite were log-transformed to achieve homoscedasticity and normality of residuals. Using the nlme package (version 3.1) for linear mixed effects models, we modeled parasite prevalence within a population at a given time with a linear mixed effects model that included *Epichloë* inoculation treatment and a third order polynomial of days after infection, as well as their interaction, as predictors (Pinheiro, Bates, DebRoy, Sarkar, & Team, 2013). We determined the appropriate polynomial to utilize using AIC. We included random slopes to account for repeated measures of the populations.

We additionally evaluated if *Epichloë* inoculation treatment affected peak parasite prevalence. We quantified peak parasite prevalence as the highest proportion of leaves infected with the parasite at a given time point at the population-level. We tested if the *Epichloë* inoculation treatment affected peak parasite prevalence with a linear model that included *Epichloë* inoculation treatment as the predictor.

Under density-dependent transmission, the contact rate between susceptible and infected individuals depends on the host population density; transmission rate therefore increases with density. While host population density (here, the number of leaves in a population) was not measured in the 2018 experiment, aboveground biomass was measured at the conclusion of the experiment. To investigate if host population density significantly correlated with biomass, we used data from the 2017 field mesocosm experiment in which host density was measured explicitly. We used a ranged major axis regression model (implemented in lmodel2, version 1.7, (Legendre & Oksanen, 2018)) to investigate if there was a correlation between total aboveground biomass of plants and the total number of leaves in a given population at the end of the experiment. Both biomass and total leaves were log-transformed. Based on that correlation, we then used aboveground biomass as a proxy for host density of each population. Specifically, to test if the effect of the *Epichloë* inoculation treatment on parasite peak prevalence was due to variation in host density, we added total aboveground biomass to the model as a covariate.

## Results

*Epichloë*-inoculated populations tended to experience more disease than *Epichloë*-free populations. *Epichloë*-inoculated populations experienced 8.3% higher disease integrated over time as measured by AUDPS calculated with disease prevalence (Figure 1, Table 1, p = 0.051). Results when AUDPS was calculated with disease severity were generally consistent, although statistical support was weaker (Supplementary results; Table S3, Figure S4, S5). A mixed model of prevalence over time complemented these results, as the *Epichloë* inoculation treatment tended to increase parasite prevalence over time, though this finding had weak statistical support (Figure 2A, Table 2, p=0.07). The *Epichloë* inoculation treatment did clearly increase the peak parasite prevalence. Specifically, the *Epichloë*-inoculated populations had a 27% higher peak prevalence than *Epichloë*-free populations, with mean peak parasite prevalences of 0.43 and 0.34, respectively (Figure 2B, Table 3, p=0.007). Together, these results suggest that while this endophyte may impact spread of a parasite across a population over time, it most clearly leads to higher parasite prevalence at the peak of an epidemic.

**Figure 1:**
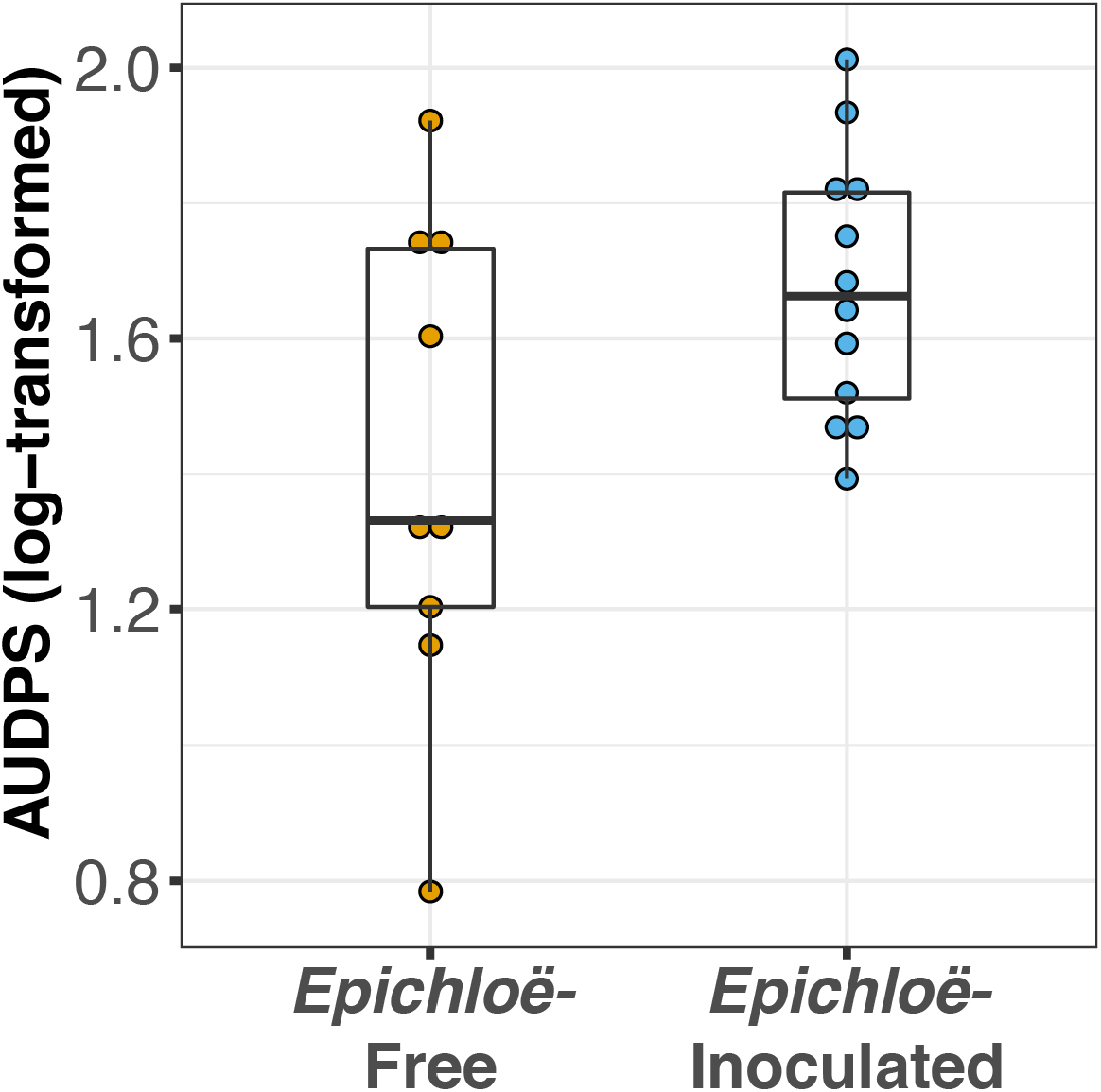
*Epichloë*-inoculated populations had greater disease prevalence integrated over time. Disease prevalence was integrated over time for each population as log-transformed AUDPS (area under the disease progress stairs). AUDPS was estimated using disease prevalence over the eight surveys until each population’s peak. Boxplots show median, 25^th^, and 75^th^ percentile, with dots showing data for each population replicate. Whiskers extend to the lowest and highest values no further than ± 1.5 times the inter-quartile range.

**Table 1:** *Epichloë*-inoculated populations tended to have heavier disease prevalence integrated over time as log-transformed AUDPS, though this difference had weak statistical support.

|  | AUDPS: Parasite Prevalence |  |  |  |
| --- | --- | --- | --- | --- |
|  | F | numDF | denDF | P |
| *Epichloë*<br>Inoculation | 4.33 | 1 | 20 | 0.051 |

**Figure 2:**
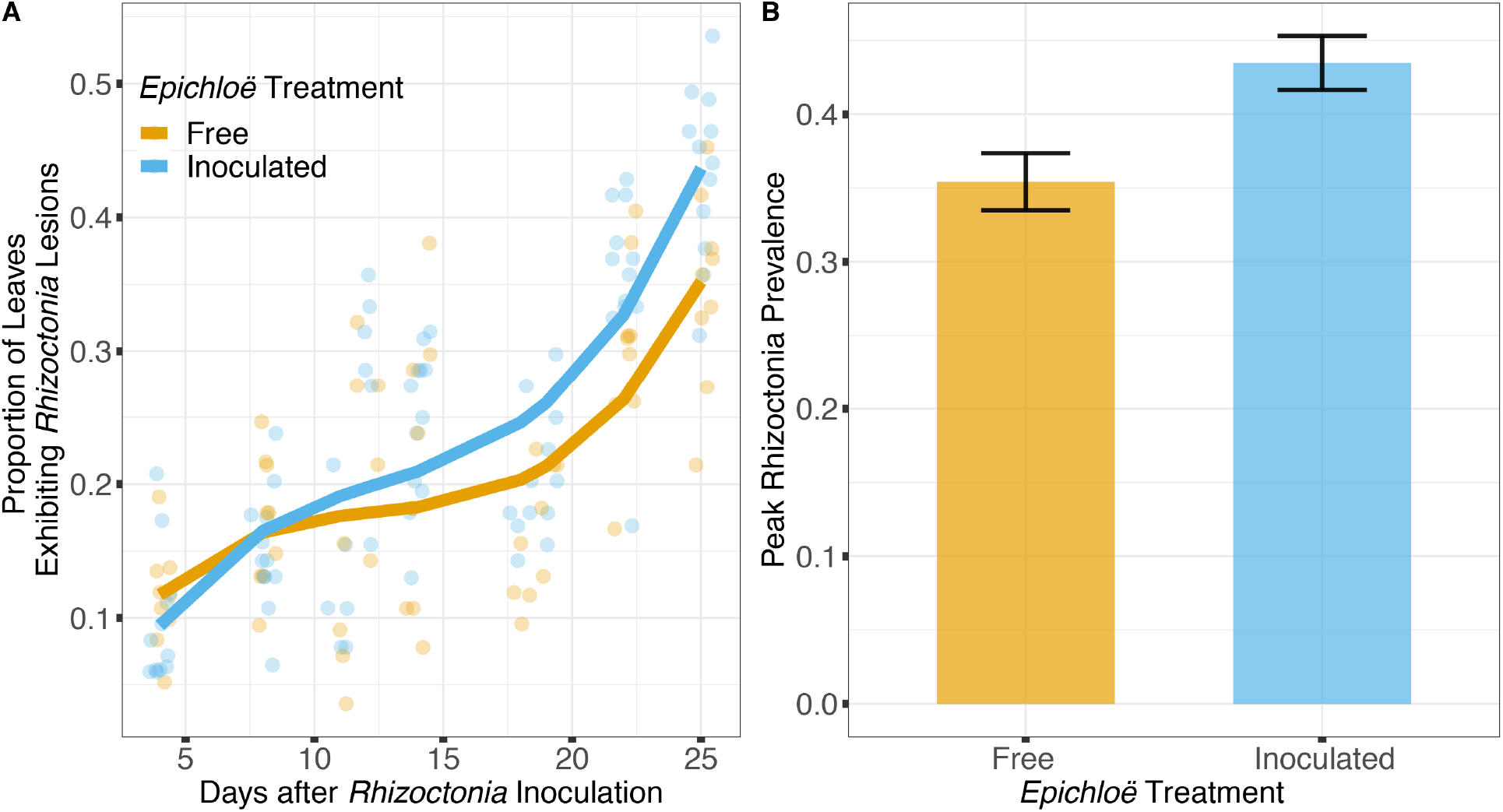
*Epichloë*-inoculated populations tended to have greater disease prevalence over time, though this difference had weak statistical support in a linear mixed model. More clearly, *Epichloë* inoculation increased peak parasite prevalence experienced by a population. **A**. Bold lines are model-predicted means of parasite prevalence over the course of 4 weeks post-inoculation with the parasite for populations in which *Epichloë* was inoculated (blue) and populations in which *Epichloë* was absent (orange). **B**. *Epichloë*-inoculated populations had 8.6% higher peak parasite prevalence peak prevalence than *Epichloë*-free populations. Bars indicate mean peak parasite prevalence and error bars are ± 1 SE.

**Table 2:**
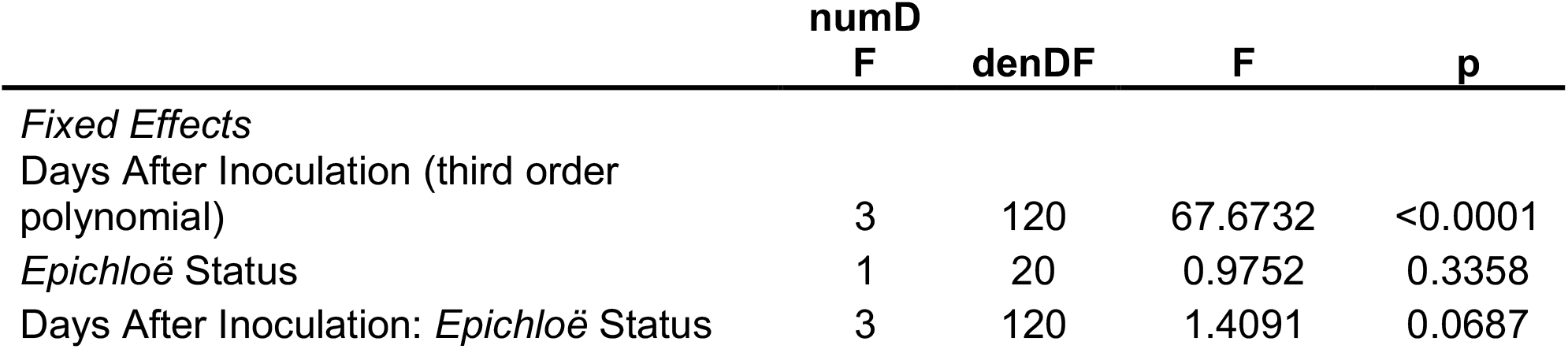
*Epichloë*-inoculated populations tended to have greater disease prevalence tracked over time in a linear mixed model, though this difference had weak statistical support.

**Table 3:** *Epichloë* inoculation clearly increased peak parasite prevalence experienced by a population.

|  | numDF | denDF | F | p |
| --- | --- | --- | --- | --- |
| *Epichloë* inoculation | 1 | 20 | 9.218 | 0.006795 |

In addition to impacting parasite prevalence and disease severity, this endophyte also impacted host population aboveground biomass. Specifically, populations with higher prevalence of *Epichloë* also had higher aboveground biomass (Figure 3, Table 4, p = 0.047). Motivated by the expectation that transmission of the parasite is density dependent, we investigated if higher biomass correlated with higher numbers of leaves in a population (i.*e*. host population density). As counting all leaves was not feasible in the 2018 experiment, we used data from the 2017 experiment, in which we counted all leaves in each population. In the data from 2017, we investigated if there was a correlation between the aboveground biomass of the plant population measured at the end of the experiment and the total number of leaves surveyed at the end of the experiment. Population-level aboveground biomass and the number of leaves in a population were clearly positively correlated in 2017 (Figure 4, Table 5, Marginal R^2^=0.145, p = 0.045). If we assume that the correlation between population biomass and leaf number from 2017 also held in 2018, then that suggests that in analyzing the 2018 experiment, host population aboveground biomass can be used a proxy for host population density.

**Figure 3:**
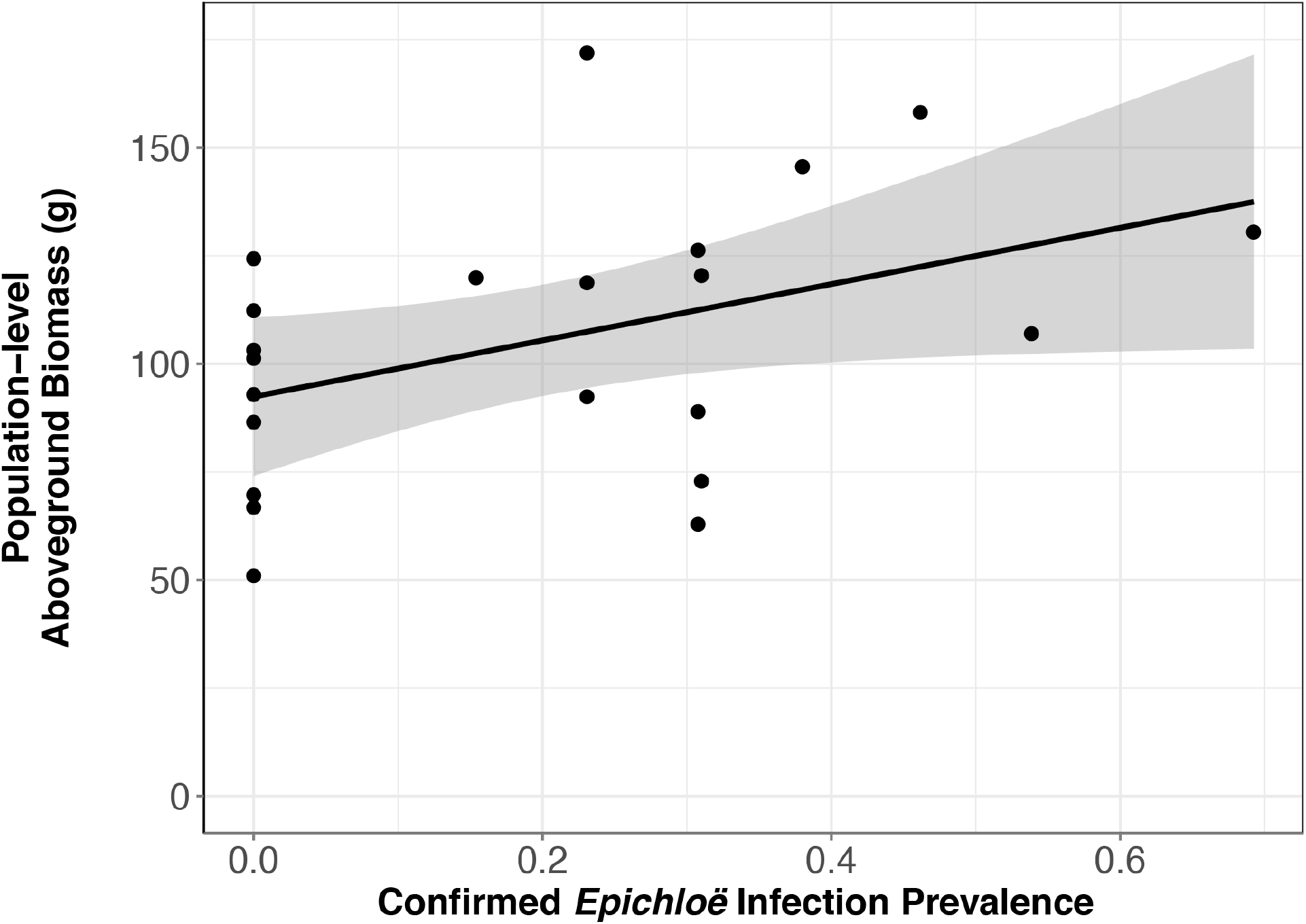
*Epichloë* infection prevalence was associated with greater host aboveground biomass at the population level. Each point represents a host population, and the bold line represents the best fit linear model. At the population level, confirmed *Epichloë* prevalence was clearly positively associated with population-level host aboveground biomass at the end of the experiment (p = 0.047).

**Table 4:** *Epichloë* prevalence was associated with greater final aboveground biomass of the host population.

|  | numDF | denDF | F | p |
| --- | --- | --- | --- | --- |
| *Epichloë* prevalence | 2 | 14 | 4.465 | 0.047 |

**Figure 4:**
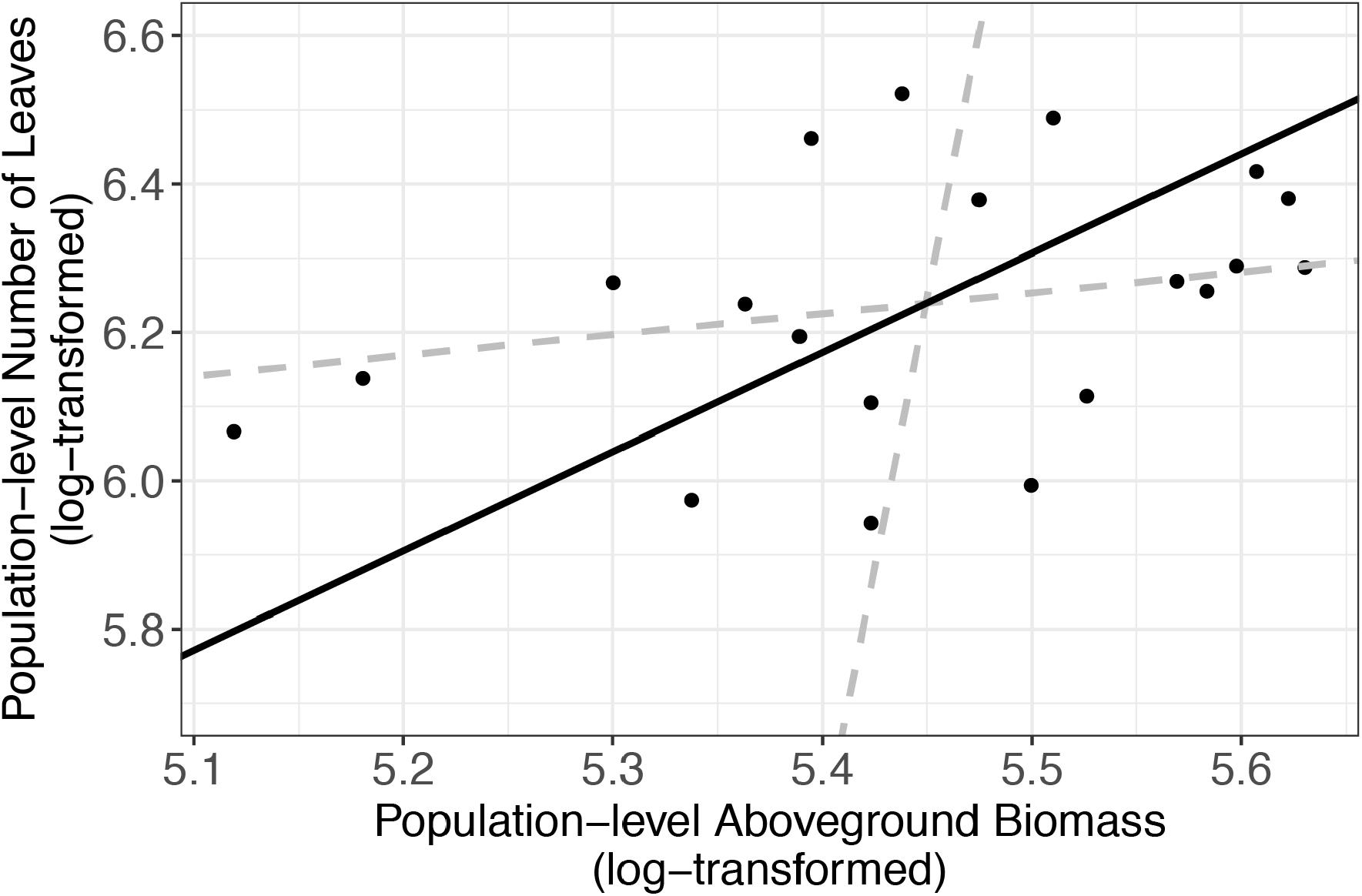
Number of leaves in a population was correlated with dry aboveground biomass of a population. We summed the total number of leaves and the biomass (in grams) of each population (the level at which other analyses were performed). Each point represents a host population in the 2017 experiment. The black solid line indicates ranged major axis fit and the gray dashed lines indicate 95% confidence intervals. The total number of leaves (log-transformed) at the end of the experiment was correlated with log-transformed dry aboveground biomass (R^2^=0.145, permutation test, p=0.045)

**Table 5:** Aboveground biomass was correlated with number of leaves at the population level (permutation test of ranged major axis regression)

|  | Intercept | Slope | Angle | P |
| --- | --- | --- | --- | --- |
| Biomass | -1.05 | 1.33 | 53.2 | 0.045 |

We then investigated whether higher levels of disease experienced by *Epichloë*-inoculated populations were driven by higher host densities. Based on the analysis of data from 2017, we used population-level aboveground biomass as a proxy for host population density and tested if biomass accounted for the effect of *Epichloë* treatment on peak prevalence. Biomass and *Epichloë* treatment explained approximately 56% of variance in peak parasite prevalence. Biomass was positively correlated with peak parasite prevalence (p=0.006), and independent of this association (interaction p=0.87), *Epichloë* inoculation increased peak prevalence (p=0.01, Figure 5, Table 6).

**Figure 5:**
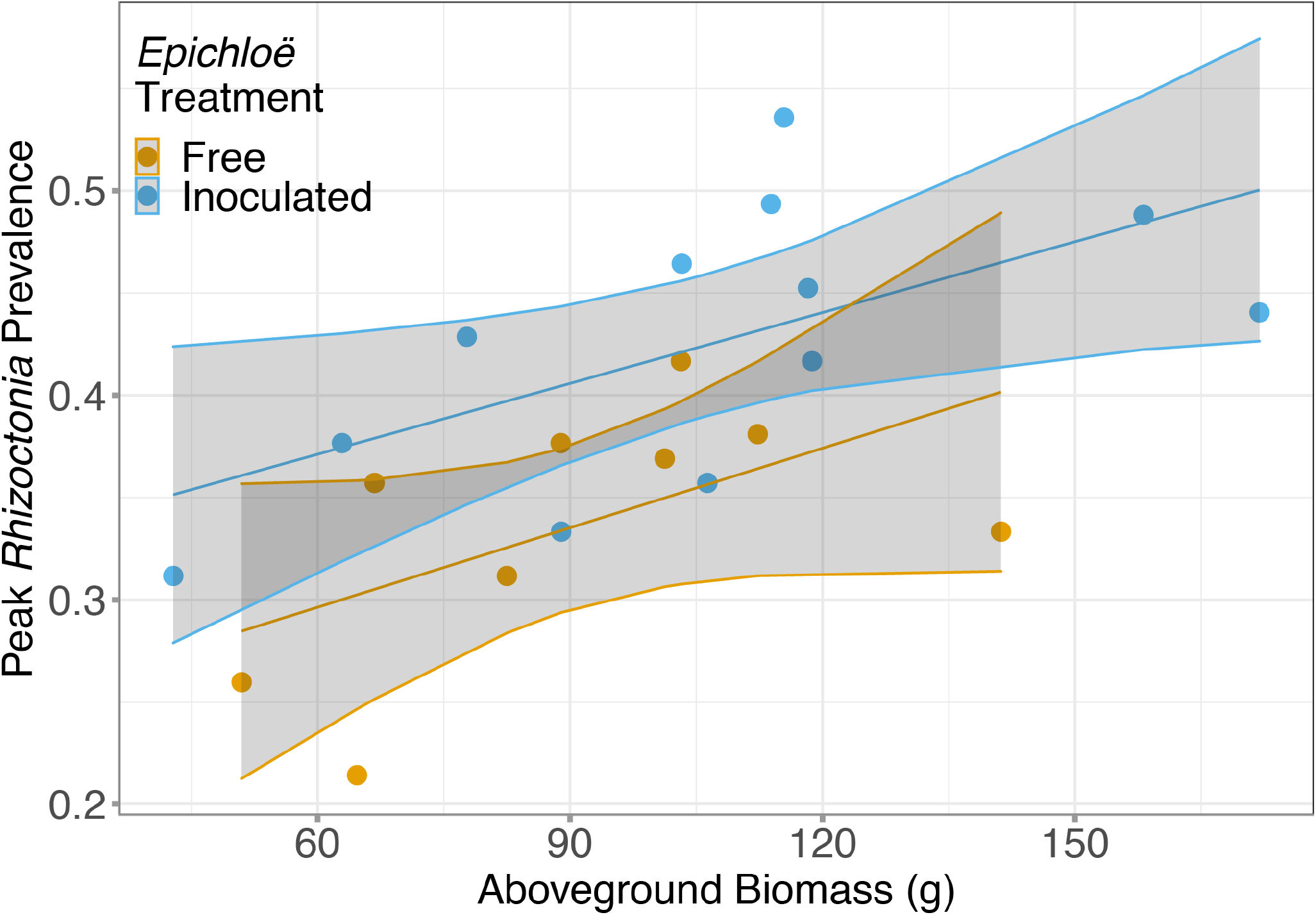
Peak *Rhizoctonia* prevalence was correlated with dry aboveground biomass. Each population (represented with a point) was either inoculated with *Epichloë* (blue) or free of *Epichloë* (orange). Biomass predicted peak parasite prevalence (p=0.006), and independent of this effect, *Epichloë* inoculation increased peak prevalence by 0.07 (p=0.01).

**Table 6:** There was a clear effect of *Epichloë* inoculation on peak prevalence, even after accounting for the association of biomass with prevalence.

|  | numDF | denDF | F | p |
| --- | --- | --- | --- | --- |
| *Epichloë*<br>inoculation | 1 | 20 | 12.6881 | 0.01 |
| Biomass | 1 | 20 | 9.126 | 0.006 |
| *Epichloë*:Biomass | 1 | 20 | 0.02 | 0.87 |

## Discussion

Our study provides experimental evidence that population-level disease dynamics can be impacted by the presence of a mutualist. Specifically, we investigated the responses of parasite spread through a host population to the presence of a mutualistic systemic endophyte under field conditions. We found that populations of tall fescue inoculated with the endophyte, *E. coenophiala*, counterintuitively experienced a higher peak prevalence of parasite, *R. solani*, over the course of the experimental epidemic.

The mutualistic relationship between cool-season grasses and vertically-transmitted systemic fungal endophytes related to *Epichloë* has been studied extensively. While *Epichloë* endophyte infection has been shown to benefit host plants by increasing resistance to herbivores and seed predators, as well as providing protection against abiotic stressors (K. Saikkonen, Faeth, Helander, & Sullivan, 1998; Kauppinen, Saikkonen, Helander, Pirttilä, & Wäli, 2016), evidence for defending against infectious disease is less consistent (Potter, 1980, 1982; Liu et al., 2006; Kari Saikkonen et al., 2013). In this experiment, we found that infection with *E. coenophiala* resulted in an increase in aboveground biomass, but there was no support for *E. coenophiala* limiting disease progression. *Epichloë*-inoculated populations actually experienced *R. solani* epidemics with higher peak prevalences. Our results are consistent with studies reporting no effect (Burpee & Bouton, 1993; Halliday et al., 2017; O’Keeffe et al., 2021) or a facilitative effect of the endophyte on a parasite (Wäli, Helander, Nissinen, & Saikkonen, 2006; Halliday et al., 2017). Vertically transmitted fungal endophytes can impact fungal parasites via resource competition and changes in host immunity, which depends on parasite feeding strategies (Kari Saikkonen et al., 2013). While we expected that *R. solani* would be inhibited by *E. coenophiala* given previous findings that *E. coenophiala* most often suppressed disease caused by a relative of *R. solani* (Pańka et al., 2013), the direction of the effect of these endophytes on parasites likely depends on host genotype and environmental conditions (Krauss & Power, 2007; Pańka et al., 2013). Further experimentation is needed to determine the mechanism underlying the potential facilitation of *R. solani* by *E. coenophiala*.

*Epichloë*-inoculated populations tended to have greater disease prevalence over time, though this difference had weak statistical support. More clearly, *Epichloë* inoculation increased peak parasite prevalence experienced by a population. We hypothesized that under density-dependent transmission, the benefit to the host individual of increased aboveground biomass, which in this case, correlated with host density (here, number of leaves), may result in a higher contact rate between hosts and consequently, higher parasite peak prevalence. While biomass and peak parasite prevalence were significantly positively correlated, consistent with density-dependent transmission, our results suggest that *Epichloë* impacted peak parasite prevalence beyond effects of biomass. One possible explanation is that *Epichloë* infection may have altered growth of *Rhizoctonia* or other processes within host individuals that scaled up to the observe effect on peak prevalence in the host population. Alternatively, given that there is evidence that parasites can impact their host’s biomass (Cordovez et al., 2017; Preston & Sauer, 2020), directly accounting for host density in models (rather than with biomass as a proxy) may more completely account for the impact of *Epichloë* on peak parasite prevalence.

There is growing evidence that the direction and magnitude of the consequences of within-host interactions are strongly affected by environmental context (Leung et al., 2018; Tracy, Weil, & Harvell, 2018). Our study provides a contribution to this understanding, as it expands upon previous work investigating the interaction between this hypothesized mutualist and this parasite under controlled settings (O’Keeffe et al., 2021) by interrogating the impacts of the same interaction on parasite spread at the population level under field conditions. Foliar fungal parasites have been studied extensively and can serve as a suitable model system to investigate microbiome/parasite interactions under field settings. Here, we show that field mesocosm experiments offer the ability to investigate the effect of within-host microbial interactions on parasite spread.

Within-host microbial interactions can influence natural epidemics in complex ways (Mordecai, Gross, & Mitchell, 2015; P. A. Clay, Cortez, Duffy, & Rudolf, 2019). Across hosts, we found that populations inoculated with a mutualist counterintuitively experienced higher peak prevalence of this parasite. These results suggest that within-host interactions among parasites and non-pathogenic microbes can impact epidemic dynamics, and we propose that field mesocosm experiments can yield important insight into disease dynamics across populations under field settings.

## Conflicts

The authors declare that the research was conducted in the absence of any commercial or financial relationships that could be construed as a potential conflict of interest.

## Funding

This work was supported by the NSF-USDA joint program in Ecology and Evolution of Infectious Diseases (USDA-NIFA AFRI grant 2016-67013-25762). K.R.O. was supported by a graduate research fellowship from the National Science Foundation.

## Acknowledgements

We thank Marc Cubeta for helpful conversations about the biology of Rhizoctonia, James Umbanhowar for help with statistics, and Fletcher Halliday, Ignazio Carbone, Corbin Jones, and Elizabeth Shank for helpful comments on experimental design and analyses. We also thank Brooklynn Joyner, Charlie Muirhead, Storm Crews, and Julie Long for help in the field and Tim Phillips and Rebecca McCulley for the tall fescue seed.

## Data Accessibility Statement

The datasets generated and analyzed for this study can be found in this github repository: https://github.com/krokeeffe12/OKeeffe_fieldmesocosm

## Notes

https://github.com/krokeeffe12/OKeeffe_fieldmesocosm

### Competing Interest Statement

The authors have declared no competing interest.

