## Supplementary Material for "A microbial mutualist within host individuals increases parasite transmission between host individuals: Evidence from a field mesocosm experiment"

Supplement

### **<u>Supplementary methods</u>**

#### **2017 Field Experiment**

<u>Experimental Design and Setup</u>

We investigated how within-host microbial interactions impact parasite transmission by conducting field mesocosm experiments in 2017 and 2018. The experimental design was largely the same between the two years (with the 2018 methods described in full in the main text). Here, we report only the differences in 2017.

To test how the endophyte *Epichloë* affects parasite spread across a host population, we manipulated *Epichloë* presence in populations. We planted a total of 20 populations, 10 *Epichloë*-inoculated and 10 *Epichloë*-free. Two randomly selected populations in each endophyte treatment (4 total) were not inoculated with the parasite and served as experimental controls.

The 260 plants within the experiment were propagated from *Epichloë*-free or *Epichloë*-inoculated seed produced by Tim Philips at the University of Kentucky and the Noble Research Institute in Ardmore Oklahoma, respectively. Seed was germinated on June 21, 2017 and grown in a greenhouse for 6 weeks. All plants except for the central plants were transplanted into the contained field mesocosm experiment on Monday, August 2, 2017. Populations consisted of plants all belonging to the same endophyte category (all *Epichloë*-inoculated or *Epichloë*-free). Notably, in contrast to the 2018 experiment, central plants in the 2017 experiment were not all *Epichloë*-free. Instead, central plants belonged to the same category as the population in which it was transplanted. The populations were fully randomized in a 2 x 10 layout, with narrow paths separating populations (Figure S1). The plants were given two days to acclimate to the field prior to the introduction of the parasite.

The plants that would ultimately be planted in the central position of the populations were transferred to growth chambers on August 2, 2017 and inoculated with the parasite or mock-inoculated. After two days, all plants inoculated with *R. solani* exhibited parasite symptoms and were transplanted into the field mesocosm experiment on August 4, 2017.

Data Collection

Following the placement of the central inoculated plant, at varying intervals over 72 days, twice a week, all leaves on every plant were selected and observed for the presence or absence of damage caused by *R. solani*, as well as any other damage. Specifically, leaves were surveyed 3, 4, 5, 6, 7, 10, 12, 18, 25, 31, 51, 58, 65, and 72 days after parasite inoculation (14 surveys total). No severity surveys were conducted.

Data Analysis

*Epichloë*-inoculated seed did not always result in *Epichloë* detection in aboveground tissue. Overall, we detected the *Epichloë* in aboveground tissue in 49.5% of endophyte-inoculated plants. This resulted in variation in *Epichloë* prevalence among the *Epichloë*-inoculated populations (minimum: 39.7%; maximum: 69.2%). We analyzed our data with endophyte treatment (2 levels: *Epichloë*-free or *Epichloë*-inoculated) as a predictor.

Control populations (in which the central plant was mock-inoculated) did not exhibit symptoms of the parasite, confirming that containment of populations limited environmental sources of inoculum. These populations were therefore excluded from analyses.

### Supplementary results

2018 experiment disease severity analyses:

When AUDPS was estimated with disease severity data, *Epichloë*-inoculated populations had 52% higher disease intensity over time on average than *Epichloë*-free populations, though this difference had weak statistical support (Figure S4, Table S3, p = 0.065). While the average disease intensity was much higher in *Epichloë*-free populations than *Epichloë*-free populations (52% higher), the lack of statistical support compared to that of AUDPS analyses when estimated with prevalence data is likely due to high amounts of variation within treatments. This high amount of variation may be due to limited sample size at each sampling point and the analysis was also limited by only surveying severity three times over the course of the experiment (Figure S5).

**Supplementary tables**

**Table S1**: **In 2017, *Epichloë* inoculation did not clearly impact disease prevalence over time.**

| *Fixed Effects* | numDF | denDF | F | p |
|---|---|---|---|---|
| Days After Inoculation ($4^{th}$ order polynomial) | 4 | 134 | 24.04 | < 0.0001 |
| *Epichloë* Treatment | 1 | 14 | 1.16 | 0.2988 |
| DAI:*Epichloë* Treatment | 4 | 134 | 1.49 | 0.2070 |

**Table S2: In 2018, when including all survey data, *Epichloë* inoculation did not clearly impact disease prevalence over time.**

| *Fixed Effects* | numDF | denDF | F | p |
|---|---|---|---|---|
| Days After Inoculation ($4^{th}$ order polynomial) | 4 | 139 | 37.4877 | < 0.0001 |
| *Epichloë* Treatment | 1 | 19 | 1.9288 | 0.1810 |
| DAI:*Epichloë* Treatment | 4 | 139 | 1.3121 | 0.2685 |

**Table S3: In 2018, *Epichloë*-inoculated populations tended to have heavier disease over course of experiment (when estimated with disease severity), on average, though this difference had weak statistical support.**

| | **AUDPS: Disease Severity** | | | |
|---|---|---|---|---|
| | F | numDF | denDF | P |
| *Epichloë* Inoculation | 3.85 | 1 | 20 | 0.065 |

**Supplementary figures**

**Figure S1: Prevalence over time was higher in 2018 than 2017.** Bold lines are model-predicted means of parasite prevalence, measured as proportion of infected leaves, over the course of 4 weeks post-inoculation with the parasite for populations in which endophyte was inoculated (blue) and populations in which endophyte was absent (orange). The 2017 results (dashed lines) and 2018 results (solid lines) were modelled independently.

**Figure S2: 2018 Field Mesocosm Experimental Layout.** A. Plant locations within each population. Each population was comprised of 13 plants, one plant in the center that was inoculated with the parasite with 12 plants surrounding it. The surrounding plants were at three different distances away from the central plant (12 cm, 24 cm, and

36 cm). Distance away from the central plant is shown in gray scale. Distances were such that neighboring plants had leaves in contact for most of the experiment. B. Arrangement of 26 populations.

**Figure S3: *Epichloë* inoculation did not affect parasite spread over time when including all data from 2018 surveys.** Bold lines are model-predicted means of parasite prevalence, measured as proportion of infected leaves, over the course of 4 weeks post-inoculation with the parasite for populations in which *Epichloë* was inoculated (blue) and populations in which endophyte was absent (orange).

**Figure S4: In 2018, *Epichloë* -inoculated populations tended to have heavier disease integrated over time.** Disease intensity over time of each population was calculated as AUDPS. AUDPS was estimated with disease severity over time (3 surveys). Boxplots show median, $25^{th}$, and $75^{th}$ percentile, with dots showing data for each population replicate. Whiskers extend to the lowest and highest values no further than ± 1.5 times the inter-quartile range.

**Figure S5: 2018 field experiment severity estimates over time.** Each line represents a given population colored by treatment (*Epichloë*-inoculated is blue; *Epichloë*-free is orange). The average disease severity (as estimated by percent leaf damaged) was estimated once per week for three weeks.

Figure S1

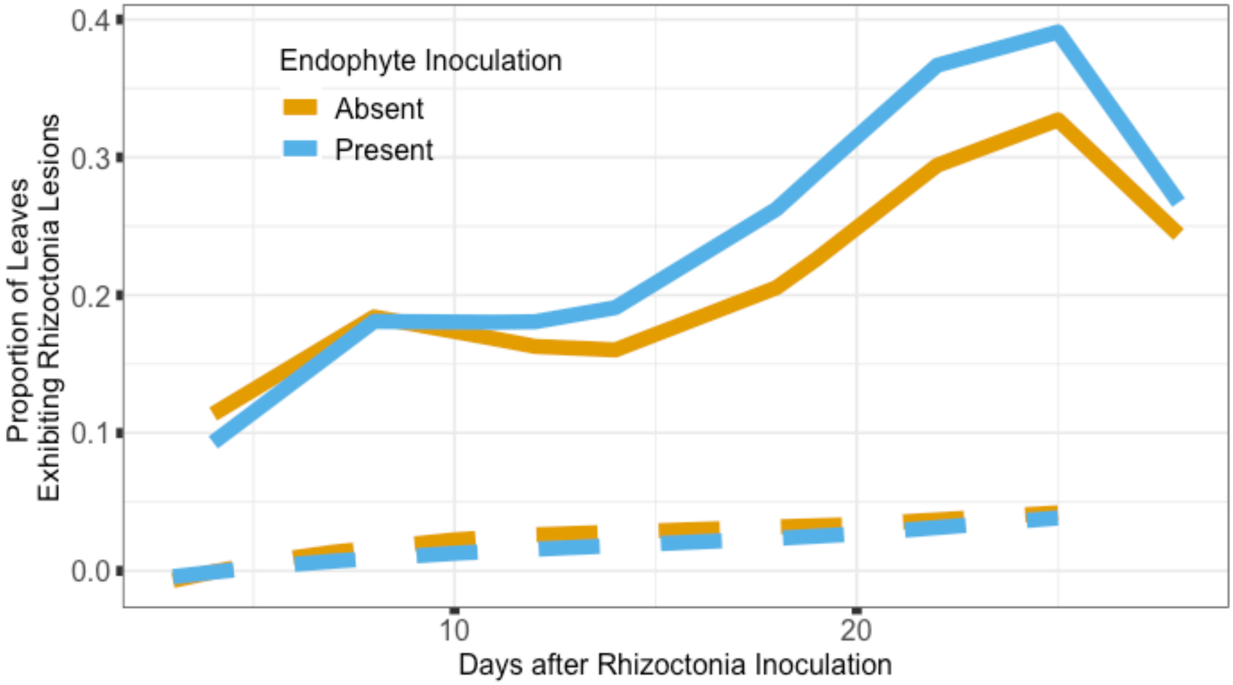
Endophyte Inoculation
Absent
Present
Proportion of Leaves
Exhibiting Rhizoctonia Lesions
0.4
0.3
0.2
0.1
0.0
10
20
Days after Rhizoctonia Inoculation

Figure S2

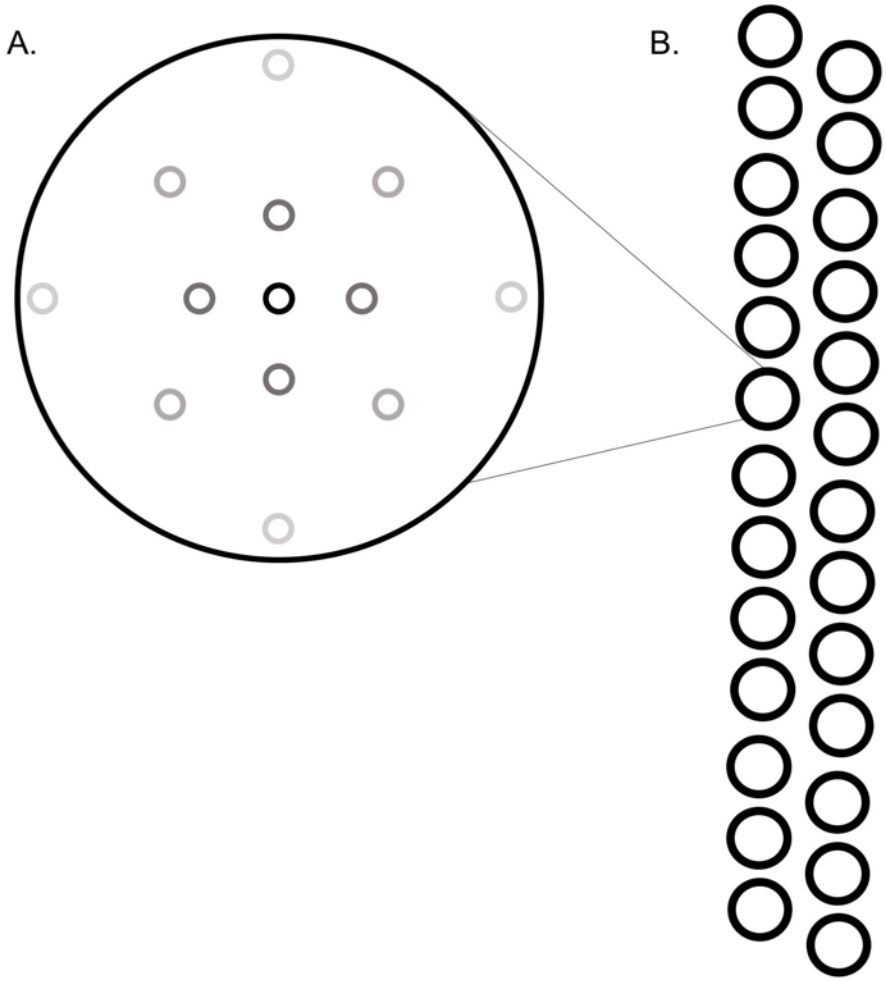

Figure S3

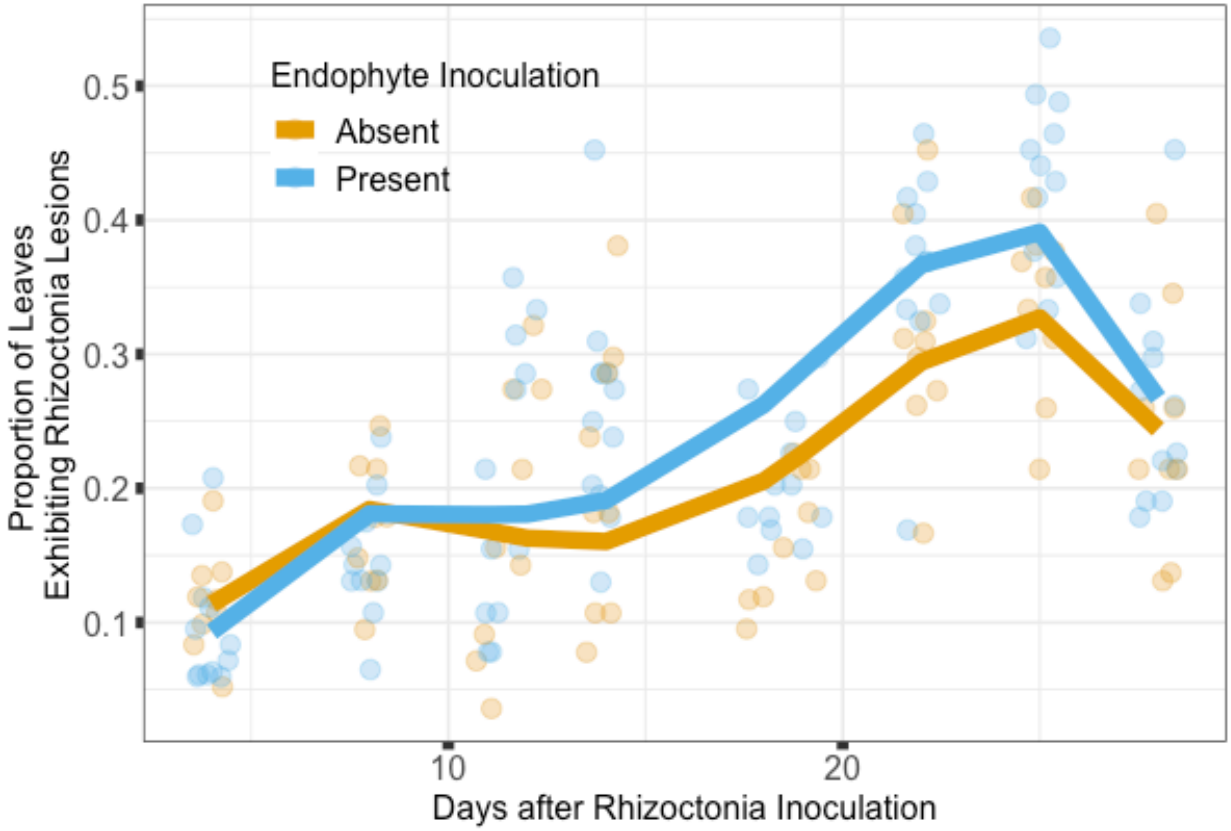
Endophyte Inoculation
Absent
Present
Proportion of Leaves
Exhibiting Rhizoctonia Lesions
0.1
0.2
0.3
0.4
0.5
10
20
Days after Rhizoctonia Inoculation

Figure S4

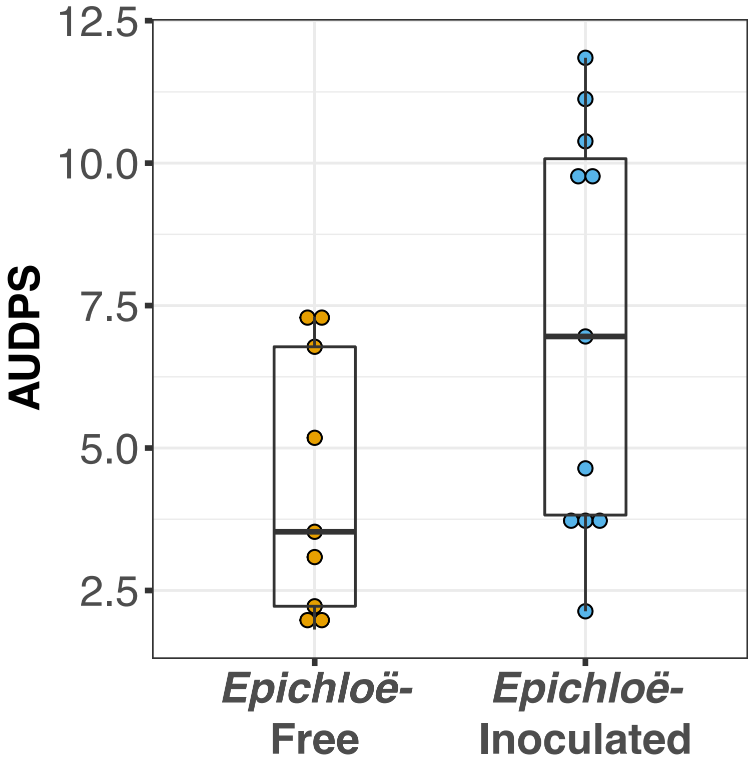

Figure S5

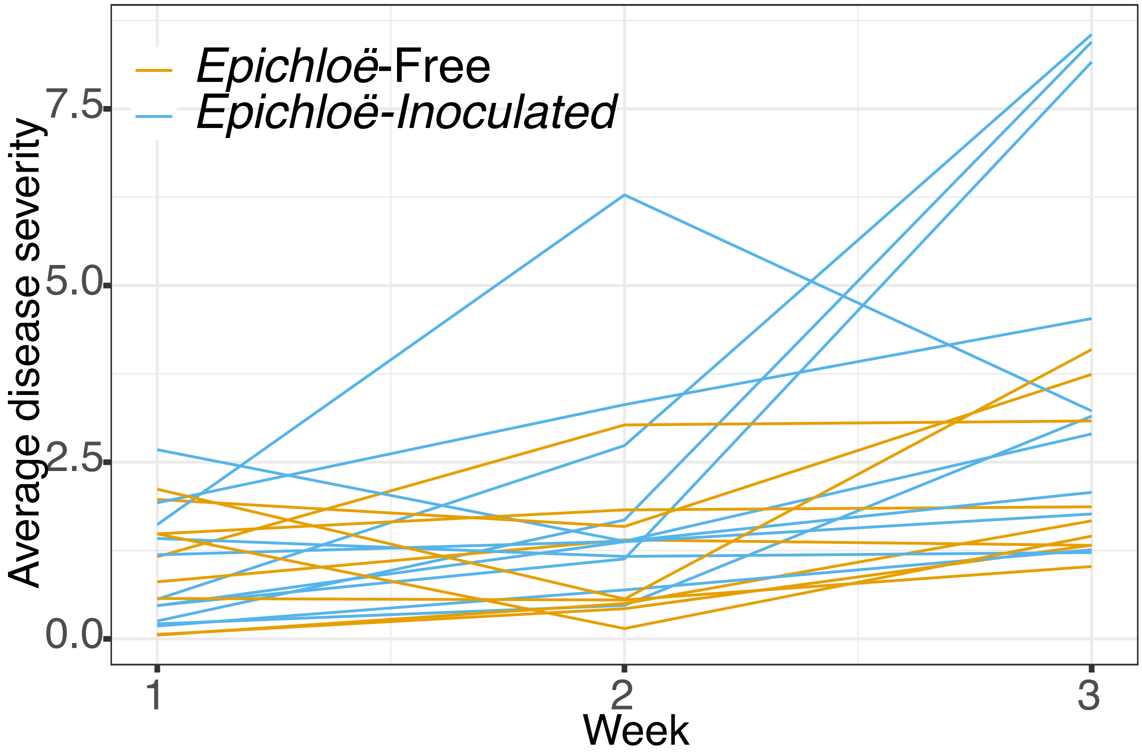

Epichloë-Free
Epichloë-Inoculated
7.5
5.0
2.5
0.0
Average disease severity
1
2
3
Week
